# First estimates of the population growth rate of the parasitic honey bee mite *Tropilaelaps mercedesae* in *Apis mellifera* colonies

**DOI:** 10.64898/2026.07.06.736813

**Authors:** Dan Aurell, Rogan Tokach, Bajaree Chuttong, Pichet Praphawilai, Léna Barascou, Todd D. Steury, Kate Duffy, Chuleui Jung, Hyunha Oh, Selina Bruckner, Geoffrey R. Williams

## Abstract

A parasitic mite of honey bee brood (*Tropilaelaps mercedesae*), is spreading through populations of *Apis mellifera* honey bees in new regions and poses a major threat to honey bee health. Despite its clear threat, the biology of this mite is poorly understood, with gaps on such fundamental issues as how fast its populations can grow. This leaves the beekeeping world underprepared to plan for its arrival and management. In this study, we documented the growth of *T. mercedesae* populations in untreated *A. mellifera* colonies in Thailand and South Korea, and did the same for another parasitic mite (*Varroa destructor*) when possible. We found that the population growth of *T. mercedesae* was variable but could reach high levels (daily *r* of 0.010, 0.036, and 0.057), while the population growth of *V. destructor* (*r* = 0.021) matched previous estimates. Our results indicate that *T. mercedesae* populations can grow rapidly but they do not always attain this potential. Based on our results, humidity should be studied as a potential driver of population growth. If future work can reveal key drivers of *T. mercedesae* population growth, this would help predict infestations and help design management strategies that exploit the pest’s biological vulnerabilities.

## Introduction

The Western honey bee (*Apis mellifera* L.) is the most widely managed insect pollinator worldwide. The health of this species is thus of fundamental importance to crop pollination and beekeepers’ livelihoods. Currently, two invasive parasitic mites threaten the health of this bee: *Varroa destructor* Anderson & Trueman and *Tropilaelaps mercedesae* Anderson & Morgan. Both mites feed on brood (developing honey bees), and reproduce in capped brood cells. *Varroa destructor* also feeds on adult honey bees.^1,2^ Both species of parasitic mites successfully jumped from Asian honey bee species to *A. mellifera* and have spread within *A. mellifera* populations.^3^

*V. destructor* has already spread to *A. mellifera* populations in all major beekeeping regions of the world^4^ and causes substantial damage. *T. mercedesae* is currently spreading out of its native range of South and Southeast Asia^5–8^ and has now spread both eastward to Korea and westward to Central Asia and Eastern Europe.^9^ This spread is expected to continue. Therefore, beekeepers and authorities need additional knowledge of the mite, whether they are located in the current or likely future range of *T. mercedesae*. While *T. mercedesae* is considered as great a threat to global honey bee health as *V. destructor*, it has received substantially less study than *V. destructor*.

The biology of *T. mercedesae* is poorly understood, including fundamental aspects like its rate of population growth. Its populations are described as rapidly growing to high levels in *A. mellifera* colonies,^2^ and its population growth is considered a major contributor to its threat to beekeeping. It has been repeatedly posited that *T. mercedesae* may pose an even greater threat than *V. destructor*. ^3,10,11^ This assessment is often based on an understanding that *T. mercedesae* mites’ populations grow faster than those of *V. destructor*.^3^ However, this belief is supported primarily by indirect evidence: in tropical areas with continuous brood rearing, *T. mercedesae* tends to be present in higher numbers than *V. destructor*,^12,13^ and *T. mercedesae* shows a shorter time between successive reproductive cycles.^13^ Both lines of evidence point to higher population growth rate. Despite the understanding that *T. mercedesae* has rapid population growth, quantitative estimates of its population growth rate remain lacking. This knowledge gap leaves beekeepers, bee health advisors, and regulators unprepared even as it spreads to further regions.

The lack of estimates of *T. mercedesae* population growth rate leaves us without a basis for predicting their population growth in colonies in support of designing Integrated Pest Management (IPM) programs and biosecurity interventions. For *V. destructor*, even simple population predictions are useful to assess the benefits of specific management practices against the mite. For example, Al Toufailia et al. observed that *V. destructor* populations in England underwent 5-6 doublings in a year;^14^ when this information was combined with the 46% measured efficacy of a nonchemical intervention (drone brood trapping),^15^ it led to the conclusion that an intervention like drone brood trapping with ∼50% efficacy would need to be repeated approximately 6 times during the year to achieve sustainable *Varroa* control; this is clearly not a stand-alone method for *Varroa* control.^15^ Modeling also provides a basis for drafting year-long management programs for *V. destructor* and predicting what interventions can effectively prevent mite populations from reaching damaging levels.^16^ Estimation of mite population growth may also be useful from a biosecurity planning perspective for *Tropilaelaps* mites, as elevated *T. mercedesae* infestations in colonies can promote their dispersal to other colonies.^17^ Without estimated population growth parameters for *T. mercedesae*, even simple projections of population growth are impossible. This limits our ability to prepare first estimates of the time it takes for populations to reach levels that damage colonies and promote dispersal between colonies.

Population projections can help plan for mite management, but to make decisions in light of actual infestations, regular monitoring of infestations is essential.^18^ Population dynamics of the honey bee colony and their mite parasites are linked in a complex system, and this can easily lead to *V. destructor* populations that remain below or exceed projections. Key parameters that regulate *V. destructor* population such as brood population, amount of drone brood, and resistance traits, are variable between colonies ^19–21^ and have compounding effects when acting together. Additionally, brood and adult honey bee population dynamics are influenced by variable forage and weather conditions in specific apiary locations with follow-on impacts on *V. destructor* mite populations. Weather may also act more directly on *V. destructor* mites,^22^ adding further sources of variability to population growth. Contingent events such as the accumulation of *V. destructor* mites by drift of honey bees or robbing from other infested colonies also occur.^23,24^ For *T. mercedesae*, we expect most of these sources of variation to impact population growth, and we have the added limitation that there are more knowledge gaps concerning *T. mercedesae* biology.

Before projecting *T. mercedesae* population growth, it is important to specify what levels of infestation may be concerning. For promoting dispersal to other colonies, initial evidence indicates that when colonies have higher infestations, they are more likely to show *T. mercedesae* mites exiting on bees,^17^ promoting dispersal to other colonies as in the case of *V. destructor*. When 0-2.5% of worker brood cells were infested, no mites were detected on exiting bees, while when cell infestations ranged from 11.5-37%, mites were found on exiting bees.^17^ On this basis, we surmise that cell infestations of 10% or higher promote dispersal from colonies. We have less basis to know what levels of *T. mercedesae* infestation cause damage to colonies. Nevertheless, field observations in Thailand indicate that when the infestation levels reach approximately 10-15% of worker brood cells infested, then overt damage occurred, based on visual signs that included poor brood patterns, insufficient worker honey bee population to take care of brood, deformed wings on adult workers, and extensive brood cannibalism (Aurell, personal observation). While it is likely that infestations much lower than 10% cause substantial damage and promote dispersal, concrete thresholds are currently lacking. At present, however, we can be confident in considering a 10% *T. mercedesae* infestation rate as one that involves serious risks of inter-colony mite spread and colony damage.

In this research, we estimated population growth parameters of *T. mercedesae* in *A. mellifera* colonies by assessing mite populations and fitting these to an exponential population model. Further, we explored the implications for honey bee health, and dispersal between colonies. When possible, we also assessed *V. destructor* population growth to compare the two mite species.

## Results

Across three investigations (Thailand 2023, Korea 2024, and Thailand 2025), we estimated that *T. mercedesae* mites had a daily exponential reproductive rate (*r*) of 0.057, 0.036, and 0.010, respectively (Table 1, Figure 1). The mean of these estimates (a daily *r* of 0.033) corresponds to a 2.74-fold population increase of *T. mercedesae* mites over 1 month.

**Figure 1.**
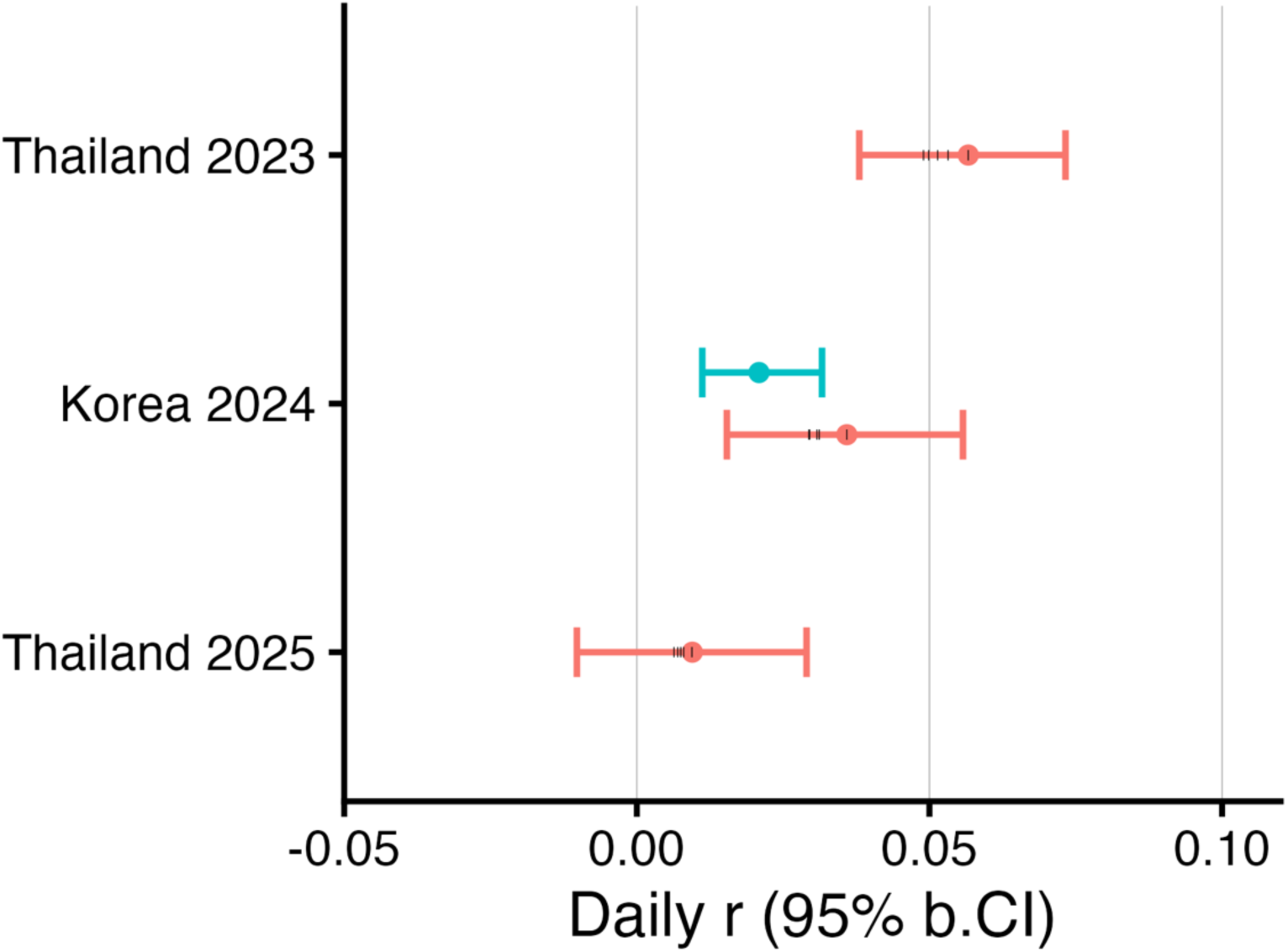
Estimated daily rate of mite population increase (*r*). Daily *r* is shown for *Tropilaelaps mercedesae* (red) and *Varroa destructor* (blue) foundress mites in *Apis mellifera* honey bee colonies across three investigations. Bootstrap means are shown and error bars denote 95% bootstrap confidence intervals. For *T. mercedesae*, when bootstrap resampling generated 0 infested cells, we inferred 0.1 foundresses / 200 cells; vertical tick marks indicate population growth rates based on inferred infestations of 0.2-0.5 foundresses / 200 cells.

**Table 1.** Overview of mite population growth observed across investigations. Population growth of *Tropilaelaps mercedesae* and *Varroa destructor* foundress mites in *Apis mellifera* honey bee colonies across three investigations. The ratio of final mite population to initial mite population in each colony was used to calculate daily *r* (daily rate of population growth) using an exponential population model. Bootstrap means and bootstrap 95% confidence intervals are presented.

| Species | Investigation, location, months | Source | n | Days | Initial mite population | Final mite population | Daily $r$ | -fold increase over 1 month |
| --- | --- | --- | --- | --- | --- | --- | --- | --- |
| *Tropilaelaps mercedesae* | Thailand 2023<br>Chiang Mai<br>Jun-Aug | Tokach et al.<sup>25</sup> | 13 | 62 | 93<br>[47, 148] | 3255<br>[1560, 5345] | 0.057<br>[0.038, 0.073] | 5.63<br>[3.19, 9.34] |
|  | Korea 2024<br>Andong<br>Jul-Sep | This paper | 16 | 61 | 70<br>[31, 114] | 670<br>[258, 1089] | 0.036<br>[0.015, 0.056] | 2.99<br>[1.60, 5.47] |
|  | Thailand 2025<br>Chiang Mai<br>Oct-Nov | This paper | 10 | 28 | 458<br>[169, 821] | 777<br>[287, 1359] | 0.010<br>[-0.010, 0.029] | 1.34<br>[0.73, 2.42] |
| *Varroa destructor* | Korea 2024<br>Andong<br>Jul-Sep | This paper | 16 | 61 | 397<br>[258, 593] | 1486<br>[911, 2056] | 0.021<br>[0.011, 0.032] | 1.89<br>[1.41, 2.63] |

In the Korea 2024 investigation, we estimated a daily *r* of 0.021 for *V. destructor*. This corresponds to a 1.89-fold population increase of *V. destructor* mites over 1 month. In this investigation, we estimated that the daily *r* of *T. mercedesae* exceeded that of *V. destructor* by 0.015 [95% b.CI: -0.005, 0.031], but since the 95% confidence interval included zero, this was not statistically significant.

If population growth is sustained at the rates indicated by our three investigations, an initial introduction of only 10 *T. mercedesae* foundresses could lead to infestations exceeding 10% of worker brood cells within 90 days (3 months) for Thailand 2023 and 142 days (4.7 months) for Korea 2024. In contrast, *T. mercedesae* infestations may not reach the 10% infestation over one year based on population growth estimates from Thailand 2025 (Figure 2). Based our experimental results from Korea, infestations of *V. destructor* could take 244 days (8 months) to reach the same infestation rate (Figure 2).

**Figure 2.**
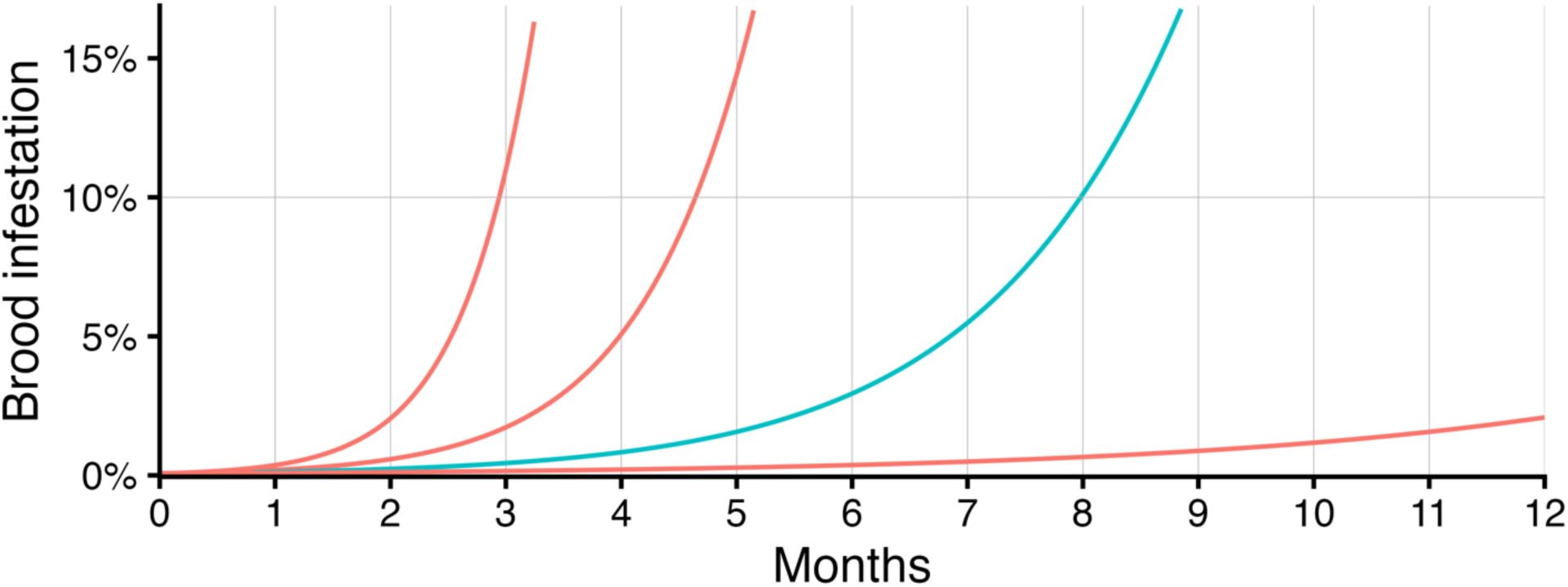
Worker brood infestation over a season based on extrapolated mite population growth. Infestation rates of *Tropilaelaps mercedesae* (red) and *Varroa destructor* (blue) foundress mite populations over 12 months in capped worker brood of *Apis mellifera* honey bee colonies. Projections were started with 10 foundress mites, and capped worker brood populations were set at a constant 15,287 cells (6 frames of worker brood, each 2/3^rd^ covered with brood). Daily rate of population increase (*r*) was obtained from three investigations: Thailand 2023; Korea 2024; and Thailand 2025. Brood infestations were corrected for multiple infestation (equation (4), see Methods section).

Rainfall during the three investigations were as follows: 7.2 mm / day, precipitation on 50/61 of days (82%) for Thailand 2023; 4.5 mm / day, precipitation on 38/54 days (70%) for Korea 2024; and 4.3 mm / day, precipitation on 19/29 days (66%) for Thailand 2025.

## Discussion

*Tropilaelaps mercedesae* already damages *A. mellifera* colonies in the regions where it exists. As its invasive range increases, it poses an increasing threat to *A. mellifera* beekeeping worldwide. Despite the urgency of responding to this pest based on an understanding of its biology, we have lacked insight into its key biological traits and parameters. In this study, we provided the first estimates of the magnitude of *T. mercedesae* population growth. These data for the first time make it possible to assess the time scale on which *T. mercedesae* infestations may rise to levels that damage honey bee colonies and promote inter-colony dispersal.

In this study we used the number of foundress mites in brood cells as a proxy for total mite population, in a study looking at population change over time. As it is one of the first studies in the honey bee research literature take this approach, we wish to critically consider the validity of the approach. For *V. destructor*, we think brood population of foundresses can be a useful proxy for total colony population but only under certain conditions. During conditions of brood rearing, *V. destructor* populations are split between the reproductive phase inside brood cells and the dispersal phase on adult honey bees, and the percentage of the total population that is in each phase can vary greatly.^23^ The ratio of brood population to adult honey bee population strongly influences the proportion of mites in both phases,^26^ and therefore can affect whether this is a good proxy for population. In the South Korea study in July-September (the only one in which we could track *T. mercedesae* and *V. destructor* mite levels, the ratio of frames of adult honey bees to frames of capped brood remained relatively stable (ratio of 2.86 at the start of the investigation; 2.04 at the end). Since this ratio remained relatively stable, we think the change in foundress mite population in brood cells was an appropriate proxy for total mite population. Corroborating the validity of our approach is that our estimated daily *r* for *V. destructor* aligned with the previous estimate of 0.021.^27^ For *T. mercedesae*, most mites are in capped brood at any time that brood rearing is ongoing, For example, studies have shown that brood is 16-20 times as infested as adult honey bees.^28,29^ Therefore, omitting mites in the dispersal phase omits a small proportion of the total mite population, and assessments of *T. mercedesae* in brood are a generally valid proxy for mite population.

This research showed that *T. mercedesae* population growth can be very rapid. Population projections based on results from two investigations (Thailand 2023, Korea 2024) indicate that after 3-4.7 months after initial infestation with a small number of *T. mercedesae* mites, the infestation of worker brood may exceed 10%, a level that is sufficient to cause acute damage to colonies (Aurell, personal observation) and that likely promotes spread to other colonies.^17^ This indicates that *T. mercedesae* could cause collapse of infested colonies and onward mite transmission to other colonies within one beekeeping season of initial infestation if these high growth rates are sustained. Nevertheless, one of our investigations (Thailand 2025) indicated a low population growth rate that we would not predict to cause damage within a one-year period.

Rapid population growth is identified as a major factor that increases the threat of *T. mercedesae* to *A. mellifera* colonies,^3^ and it has long been suspected that *T. mercedesae* has more rapid population growth than *V. destructor*^13^ based on numerous reports from the tropics of *T. mercedesae* occurring more commonly and at higher infestations than *V. destructor*.^2^ In two of our three investigations, we estimated that the daily *r* exceeded 0.021,^27^ which we use as a reference value for *V. destructor*. In our Korea 2024 investigation, we compared the population growth rate of *T. mercedesae* and *V. destructor.* We should note that the high *V. destructor* infestations in the study may have reduced *V. destructor* population growth through density-dependent effects, which would reduce the validity of a direct comparison. Nevertheless, as the estimated population growth rate of *V. destructor* matched that of previous research, we think the comparison between *V. destructor* and *T. mercedesae* is still informative. We estimated substantially higher population growth of *T. mercedesae* than of *V. destructor*. The difference in *r* was not statistically significant, but the magnitude of the estimated difference has substantial practical implications: these growth rates suggest acute colony damage from *T. mercedesae* could occur after 4.7 months, compared to 8 months for *V. destructor*, starting from low populations of both mite species. Overall, our results support the idea that *T. mercedesae* has a capacity for greater population growth than *V. destructor*.

In Korea, *V. destructor* is considered generally more prevalent than *T. mercedesae*,^30^ but there are recorded instances of *T. mercedesae* brood infestations exceeding those of *V. destructor* in July through November^30,31^ and reports of damage occurring to beekeeping operations.^8^ This variability in the relative infestations of *T. mercedesae* and *V. destructor* may be explained by the higher potential population growth rate of *T. mercedesae* overlaid with variable intensities of brood breaks, variable timing of reinfestation, variation in key factors for the mites’ population growth, and effects of beekeeper management.

We observed substantial variation in the population growth rates of *T. mercedesae* across the three investigations: the daily *r* was nearly 6 times as high in the Thailand 2023 investigation as in the Thailand 2025 investigation, and the Korea 2024 results were intermediate. Based on the very limited evidence available before we conducted this study, the view that *T. mercedesae* reproduces faster than *V. destructor* was reasonable. However, our results show that *T. mercedesae* mites do not always reach these high population growth rates. Previous research that has tracked *T. mercedesae* infestations over many months have observed both dramatic increases and also unexplained decreases in *T. mercedesae* infestation rates.^12,28,32^ These results are puzzling if *T. mercedesae* is viewed solely as a fast-reproducing organism, but make sense if it is understood as an organism that has a high capacity for population growth, yet does not always attain this capacity.

Our work highlights the importance of understanding factors that promote or inhibit population growth of *T. mercedesae* mites, especially given how variable the population growth rate seemed to be across the three investigations in this study, and across seasons in longitudinal studies. If we understand drivers of population growth, we will be better able to predict *T. mercedesae* populations, and we may be able to design interventions to inhibit their population growth.

This study provides some initial clues about factors that may regulate *T. mercedesae* populations. While we have only three investigations to compare, the data point to the possibility that high humidity may promote *T. mercedesae* population growth. The Thailand 2023 investigation in which we observed the highest *T. mercedesae* population growth also had the highest mean daily precipitation and percentage of days with precipitation (7.2 mm / day, 82% of days). The intermediate population growth rate for Korea 2024 aligned with intermediate precipitation (4.5 mm / day; 70%), and the low population growth rate for Thailand 2025 aligned with slightly lower precipitation (4.3 mm / day; 19/29 days = 66%). Clearly, these measures of precipitation are indirectly correlated to humidity, especially inside the brood nest and inside cells of honey bee colonies, but this alignment with rainfall data suggests that humidity may be important for *T. mercedesae* population growth.

While our results suggest humidity as an important factor, there is a wide array of environmental and host-parasite factors could impact *T. mercedesae* population growth, but essentially none have been systematically studied for their effect on *T. mercedesae* reproduction, mortality, or population growth. We hope that this becomes a focus area for research.

In this study, the *A. mellifera* honey bee colonies in the Thailand 2023 and the Thailand 2025 investigation were sourced from the same beekeeper. We therefore think that the genetic stock of the honey bees in these two investigations were similar, and are not the explanation for the divergent results. Similarly, while genetic factors among the *T. mercedesae* population could be important, it similarly seems unrealistic that across two years in Thailand different isolates of *T. mercedesae* with drastic differences in reproductive capabilities would be present in colonies of the same beekeeping operation.

*Tropilaelaps mercedesae* depends on honey bee brood for reproduction, and previous research has suggested a link between brood availability and *T. mercedesae* populations. In longitudinal studies, periods of brood population expansion (or simply the presence of large brood populations) seemed to be associated with periods of increase in *T. mercedesae* infestation rates and populations.^28^ While increased brood availability plausibly would facilitate reproduction by *T. mercedesae*, our investigations showed a pattern that contrasts with these observations. Our Thailand 2025 investigation had the highest capped brood areas of the three investigations, yet had the lowest *T. mercedesae* population growth. This suggests that brood availability alone is not sufficient to promote *T. mercedesae* population expansion.

*Tropilaelaps mercedesae* mites have long been understood as a grave threat to honey bee health and beekeeping in the tropics. However, there is a dawning realization that challenges from *Tropilaelaps* may extend further into temperate zones than anticipated. *Tropilaelaps mercedesae* can cause colony damage even in temperate locations like South Korea^8^ and the Russian Black Sea region.^6^ Colonies in the Russian Black Sea region sustained dramatic increases of *T. mercedesae* infestations through a study conducted in 2022-23 and 53% of colonies in the study died.^6^ With the potential for *T. mercedesae* to reach high rates of population increase, it makes sense that colony damage can occur even in the shorter brood rearing seasons typical of temperate locations, if *T. mercedesae* mites are able to persist through winter or are seasonally reintroduced.

We believe the most effective barrier to *T. mercedesae* persistence in an area is the occurrence of complete brood breaks over winter since brood is their only or major food source.^2^ (Note that a recent study based on cage trials indicated they can use alternate food sources like decaying adult honey bees to survive).^33^ Colonies in Russia’s Krasnodar region were observed rearing brood through winter^6^ despite the cold winter conditions (growing zone 7,^34^ with winter temperatures reaching -12.2 to -17.8 °C). Similarly, in Andong, central South Korea, the detection of *T. mercedesae* mites in early May before the arrival of migratory colonies^31^ suggests an ability to survive cold winters that is greater than previously appreciated. Together, these observations suggest that we may tend to overstate the extent to which temperate-climate colonies become broodless in winter, and over-estimate the suppression of *T. mercedesae* provided by cold winter weather.

Even in places where cold winters cause the local extirpation of *T. mercedesae*, colonies may suffer from introductions if infested colonies are brought in from neighboring areas. Results from Korea in which *T. mercedesae* infestations were higher in migratory colonies^35^ support the possibility of annual spread by colony migration. The results of our population predictions indicate that the timing of *T. mercedesae* reinfestation would be very important: survival of a few mites through winter could set colonies up for an extreme population growth trajectory, reinfestation early in summer could lead to problems, and reinfestation in mid summer could be completely unnoticed before winter conditions suppress mite populations again.

In conclusion, our research has validated the idea that *T. mercedesae* populations have the potential to increase at high rates, and suggest they can cause serious damage to colonies ∼3-4.7 months after infestations begin if high population growth rates are sustained. Our findings of high rates of population growth confirm that mite is a great threat to *A. mellifera* honey bees. On the other hand, our findings of highly variable population growth call for further research. Through careful study of factors influencing *T. mercedesae* survival, reproduction, and mortality, we expect that useful avenues to control this serious honey bee pest can be developed.

## Methods

### Study overview

In this study, we monitored population growth of parasitic honey bee mites (*Tropilaelaps mercedesae* and *Varroa destructor*) in untreated Western honey bee (*Apis mellifera*) colonies. In three separate investigations, we assessed mite populations of foundress mites at the start and end of observation periods 28 to 62 days in duration. The first investigation occurred in Chiang Mai, Thailand over a 62-day period in June-August 2023. The second occurred in Andong, South Korea over a 61-day period in July-September 2024. The third occurred in Chiang Mai, Thailand over a 28-day period in October-November 2025. These investigations included 13, 16, and 10 untreated colonies, respectively. Only the South Korean investigation had sufficient *V. destructor* infestations for population growth assessment. Accordingly, in the two investigations in Thailand, we only assessed *T. mercedesae* population growth.

### Overview of analyses

We used the number of foundress mites in capped brood cells of each colony as a proxy for total mite population. To determine this, we collected data on the infestation rate of capped worker brood and the number of capped worker and drone cells (see *Methodological considerations* and *Field data collection*). We used a mathematical correction to account for infestation of cells by multiple foundresses (see *Correction for multiple cell infestation*). Within a bootstrap procedure, we integrated the brood population, observed infestation rates, and correction for multiple infestation to infer the number of foundress mites per colony (see *Bootstrap analysis*). The bootstrap resampling procedure also reflected both sampling error and between-colony variability. From this procedure, we generated point estimates and bootstrap 95% confidence intervals for foundress mite populations at the start and end of observation periods, and for the daily rate of population growth (daily *r*) based on an exponential growth equation. Finally, we used the estimates of daily *r* to project mite population growth over 1 month and 12 months (see *Projecting increase of mite infestations over a season*). Analyses and plotting were conducted in R^36^ using *tidyverse*^37^ and *ggplot2*^38^ packages.

### Methodological considerations

In this analysis, we made the simplifying assumption that the infestation rate of drone brood was equal to that of worker brood. For *T. mercedesae* mites this is justified as most studies have not reported a strong preference for drone brood.^13,28,39^ While *V. destructor* mites show a substantial preference for drone brood cells,^40^ the investigation where *V. destructor* population growth could be assessed (Korea 2024), occurred during the summer resource dearth, and nearly zero drone brood was recorded. Capped drone brood was observed in only 3 of 16 colonies at the start of the observation period, accounting for only 1%, 1%, and 5% of capped brood cells in these colonies; no capped drone brood was observed in any colonies at the end of the observation period. This near absence of drone brood allowed us to use the same simplifying assumption of equal infestation of worker and drone brood for *V. destructor* as for *T. mercedesae*.

We directly recorded the number of infested worker brood cells, but wished to focus on the population of foundress mites (not foundresses and daughters), as in previous modeling papers on *V. destructor*.^27^ At the final time points (when infestations were higher than initial time points), the probability of cells being multiply infested (by more than one foundress) is expected to increase. We took this into account to avoid under-estimating population growth of mites. Given the limited evidence for the distribution patterns of *T. mercedesae* foundresses in *A. mellifera* brood cells,^31^ we based our adjustment on the assumption that the presence of infestation of multiple foundresses is based on mites randomly determining whether to infest a cell, regardless of whether it is already infested by another foundress.

### Field data collection

We assessed the infestation rate of capped worker brood cells by each mite species by uncapping and inspecting 200 capped worker brood cells per colony. For Thailand 2023 and Korea 2024, we did this by finding two frames in the colony with capped brood, and uncapping 50 worker cells on both sides. For Thailand 2025, we uncapped cells on one frame (100 worker cells on both sides). For each individual uncapped cells, we recorded whether it was uninfested, infested by *T. mercedesae*, infested by *V. destructor*, or infested by both mites, in accordance with established methods for *Tropilaelaps* research.^25,39,41^ We did not directly assess whether cells were infested by multiple foundresses of *T. mercedesae* or *V. destructor*: this determination is time-consuming and can be imprecise at late brood stages because it relies on subtle color differences between adult foundresses and offspring.^13^ Instead, we used a mathematical correction (see *Correction for multiple cell infestation*) to infer the number of foundresses from the observed number of infested cells and account for infestation of cells by multiple foundresses.

Separately, we collected data on the area of capped worker brood and capped drone brood by visually assessing percent coverage of brood on each frame (Liebefeld method)^42^ and used conversion factors^42^ to convert brood area to number of cells, to determine the number of capped worker and drone cells per colony.

Additionally, we collected data on adult honey bee population at the start and end of observation periods either using “field frames of bees”^43^ or the Liebefeld method.^42^ In two investigations (Thailand 2023 and Korea 2024), we also collected midpoint measurements of capped worker brood, drone brood, and adult honey bee population.

Finally, we retrieved data on rainfall during the observation periods from Meteostat.net. From these data we were able to assess the mean daily rainfall for each investigation, and determined what proportion of days had recorded rainfall. Rainfall data were missing for a several days. These could not be included in calculations, so when needed the calculations omitted those days.

### Correction for multiple cell infestation

Through equations (1-4), we developed an equation (5) to transform a proportion of infested cells to an inferred number of foundresses. Using probability theory, we can describe the probability of finding *x* foundress mites in a cell by equation (1).^44^ This views the infestation of individual cells by foundress mites as a random process. The more specific case of zero foundresses (i.e., cell is not infested; x = 0) is reflected in equation (2), which simplifies to equation (3). As the probability of a cell being infested is the complement of the probability of the cell not being infested, equation (4) gives the probability of 1 or more foundresses (i.e., cell is infested; *x* > 0).

*P* = Probability

*n* = Number of cells inspected

*x* = Number of foundresses present in a particular cell

*k* = Number of foundress mites among all *n* inspected cells

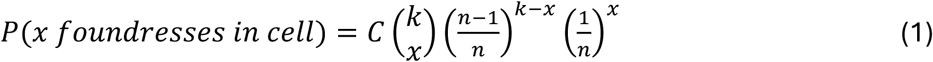

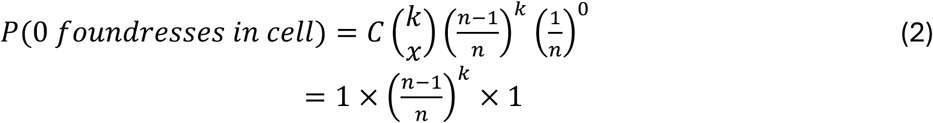

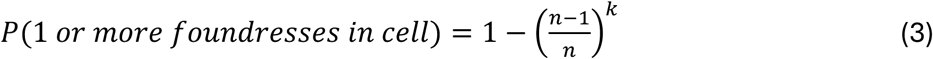

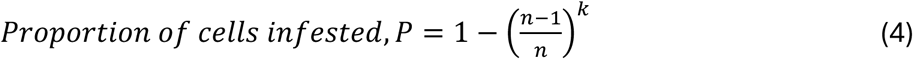

To infer *k* (number of foundress mites) that resulted in *P* (proportion of cells infested) in *n* (number of inspected cells), we can rearrange this equation for *k,* equation (5).

*P* = Proportion of cells infested

*n* = Number of cells inspected

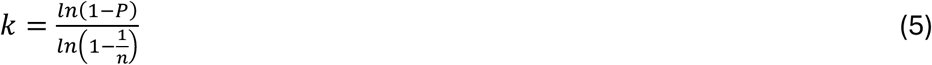

### Bootstrap analysis

We used a hierarchical bootstrap procedure^45^ to analyze separately the data from each of the three investigations. Each bootstrap iteration (10,000) included steps to reflect both the between-colony and within-colony variation (sampling error). To reflect between-colony variation, we sampled with replacement from the colonies in the investigation, generating resamples of the same number of colonies as before (i.e., 13, 16, and 10 colonies for the three investigations). Within each resampled colony, sampling error was modelled using a parametric bootstrap, in which the number of infested cells was drawn from a binomial distribution of 200 trials (cells) with infestation probability *p*, equation (6).

*c_rand_ =* Resampled count of infested cells per 200 cells

*p* = Infestation probability

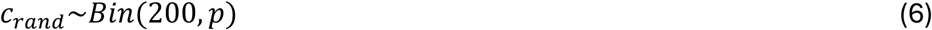

When *p* was defined as the observed proportion of infested cells, this inflated the number of zeros in the resampled data set beyond that of the original data set. This is because low observed values (e.g. 1/200 cells infested; *p* = 0.005) frequently resample to zero, whereas observed zeros cannot increase. To address this issue, we modified the infestation probability by adding a small adjustment term, equation (7). We empirically obtained the value of *adj* to ensure that the mean number of zeros per bootstrap iteration in the resampled dataset matched that of the observed dataset. For *T. mercedesae*, the value of *adj* was set as 0.00304, 0.00186, and 0.00125, for Thailand 2023, Korea 2024, and Thailand 2025, respectively. For *V. destructor* for Korea 2024, the value of *adj* was set as zero since very few zeros were generated in the resampling process due to the higher observed infestations.

*c* = Number of cells infested

*n* = Number of cells inspected

*adj* = Adjustment factor

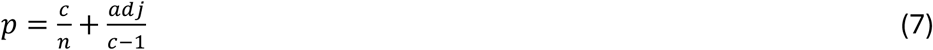

From the resampled number of infested cells in 200 cells (*c_rand_*), we used equation (5) to infer the corresponding number of foundress mites per 200 cells.

To estimate the colony-level population of *T. mercedesae* and *V. destructor* foundress mites, we multiplied the total number of capped brood cells (worker plus drone cells) by the inferred number of foundresses per 200 cells, and divided by 200. This provided an estimate of total number foundress mites of both species per colony. This is analogous to the procedure suggested by Dietemann et al.^46^ for integrating infestation rate with host population to estimate colony-level mite populations. In our case, we did not assess mite population on adult honey bees, as a small fraction of *T. mercedesae* populations are found on adult bees.^28,29^

Using the above methods, we calculated, for each colony in the bootstrap data set, the initial and final population of *T. mercedesae* and *V. destructor* foundress mites in capped brood cells. When the estimated foundress population was zero, this made calculations impossible, so these were slightly raised upwards. We tested several approaches: in the first approach, such zeros were adjusted to a population size of 1; in a second approach we inferred that the cell infestation was low, corresponding to 0.1 to 0.5 foundresses / 200 cells. Since the first approach gave rather extreme estimates (Supplemental Figure S1), we chose the second approach, and we chose to infer a mite infestation of 0.1 foundresses since we think it would be preferable to over-estimate *T. mercedesae* reproduction than to under-estimate it.

We then computed the ratio of final to initial mite population to represent “times-as-many” population growth. Based on this growth metric, we extracted the daily rate of population increase, equation (8), which corresponds to the exponential growth model used by Harris et al.^22^, but parameterized on a daily rather than weekly timescale.

*F_t_* = Final population of foundress mites (end of observation period)

*F_0_* = Initial population of foundress mites (start of observation period)

*r* = Daily rate of population increase

*t* = Number of days in the observation period

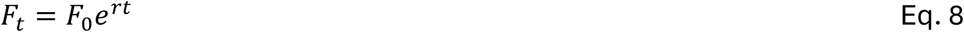

From this dataset, we summarized results using established bootstrap methods. First, for each bootstrap iteration, we determined the mean of each parameter of interest (initial mite population, final mite population, and daily growth rate *r*). Second, we computed the bootstrap estimate as the mean of these iteration-level means. To obtain bootstrap 95% confidence intervals (b.CI), we found the value of each parameter’s mean that corresponded to the 2.5% and 97.5% quantiles of the iteration means.

To set the population growth on a more interpretable scale, we also calculated the expected population growth over one month using equation (8). The fold-change in population growth was obtained using the bootstrap estimated mean of *r* and a period of 30.5 days (to reflect the length of an average month). We determined bootstrap confidence intervals for the fold-change by applying the lower and upper confidence limits of daily *r* into equation (8).

### Projecting increase of mite infestations of brood over a season

To explore the implications of the observed population growth rates, we conducted simple deterministic population projection for brood infestations in a colony over a beekeeping season. It is easier to assess the potential for a mite population to cause colony damage if it is expressed as an infestation rate rather than as a numerical mite population. We provided context by overlaying populations of mite foundresses onto a brood population that (to simplify assumptions) was kept constant through the projection. Each simulated colony had 6 frames of brood (Langstroth Deep size), each two-thirds covered with worker brood. Of the resulting 26,752 cells of brood per colony,^47^ 12 of every 21 cells were considered capped, (as 12 of 21 days of worker brood development occur in the capped phase) leading to a capped worker brood population of 15,287. This brood population was chosen to be easily relatable for beekeepers and to align with directly measured brood populations during main brood rearing seasons.^20^ Nevertheless, the projected infestations were largely driven by the increase in mite population, and varying the brood population had a limited effect on projected infestations.

We began each population projection with 10 foundress mites (*T. mercedesae* or *V. destructor*) per colony and calculated daily populations of foundress mites using equation (8) and the estimates of *r* derived from our three investigations. To obtain the infestation rate of worker brood, we used the set number of capped worker brood cells and the number of foundress mites on each day; these two numbers were put into equation (4) which provided a number of infested cells, corrected for multiple infestation. Then, the resulting number of infested cells was divided by the total number of capped cells to obtain a percentage of infested cells.

We made the following assumptions: the population growth rate remained constant throughout the projection at the mean estimated daily *r* indicated by each investigation; density-dependent effects did not reduce mite population growth when infestations became high; and mite immigration or emigration did not differ from those in the investigation.

## Acknowledgements

This research was made possible, in part, by Cooperative Agreements from the United States Department of Agriculture’s Animal and Plant Health Inspection Service (APHIS) and Agricultural Research Service (ARS): USDA-AP23PPQSCT00C067, USDA-AP24PPQSCT00C025, USDA-AP25PPQS7T00C182, USDA-58-6066-9-042, and USDA-58-6066-3-029. It was also supported by the United States Department of Agriculture’s National Institute of Food and Agriculture, under Multi-state Hatch Project NC1173. Statements in this publication may not necessarily express the views of USDA.

We also acknowledge support from the Alabama Agricultural Experiment Station, and from Project Apis m. and Bayer Healthy Hives under PROJECT APIS M-RA 382. This research was partly supported by Chiang Mai University.

## Author contributions

DA: Conceptualization, Methodology, Investigation, Formal Analysis, Visualization, Writing – Original Draft Preparation

RT: Methodology, Investigation, Writing – Review & Editing

BC: Supervision, Writing – Review & Editing

PP: Investigation, Writing – Review & Editing

LB: Investigation, Writing – Review & Editing

KD: Methodology, Writing – Review & Editing

CJ: Supervision, Writing – Review & Editing

HO: Investigation, Writing – Review & Editing

SB: Investigation, Writing – Review & Editing

GRW: Funding acquisition, Supervision, Methodology, Writing – Review & Editing

## Data availability statement

Data and analysis code will be openly provided on GitHub at the time of submission to a journal [will be replaced with Zenodo DOI upon acceptance].

## Supplemental Material

**Supplemental Figure S1.**
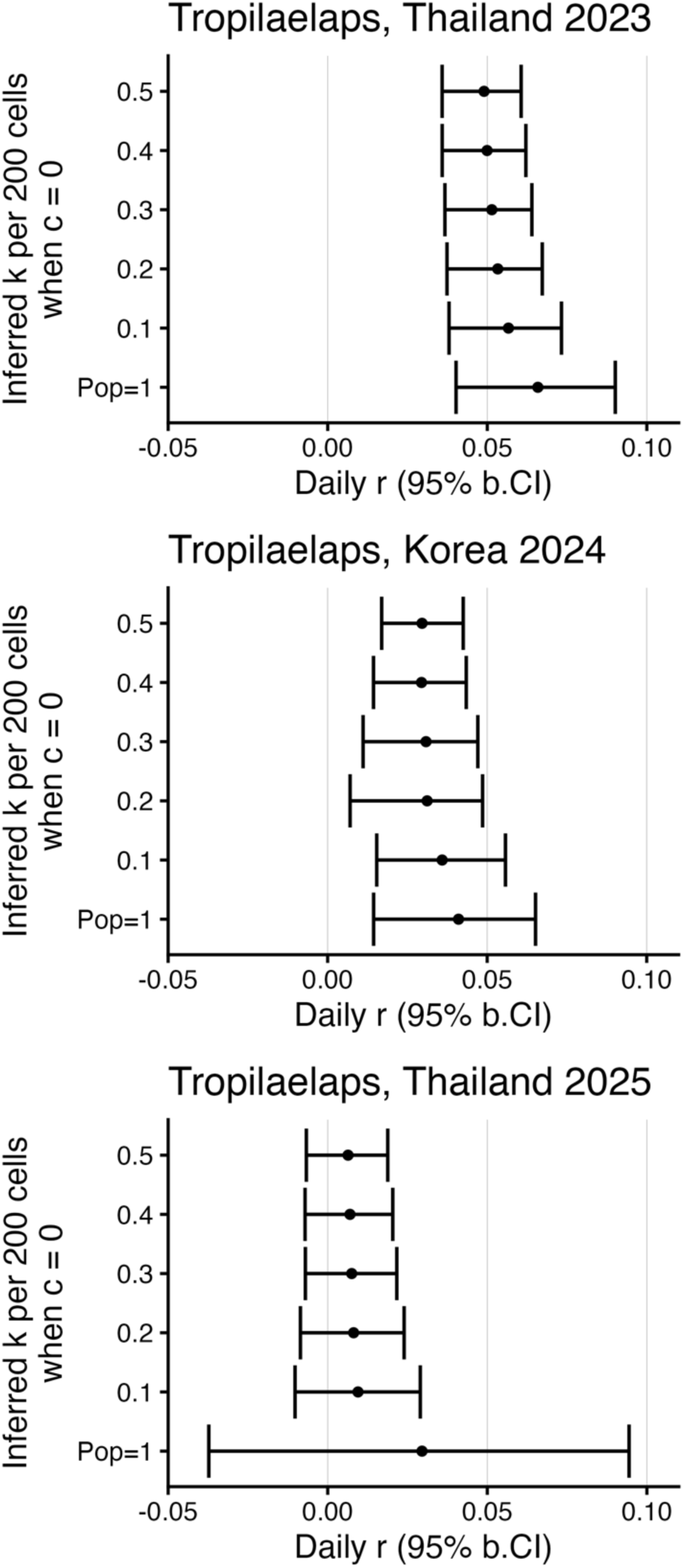
Sensitivity of *r* of different approaches to infer populations of *Tropilaelaps mercedesae* foundresses. When resampling generated 0 mites / 200 inspected cells, an adjustment was required. A *k* of 0.1-0.5 indicates that 0.1-0.5 foundresses / 200 cells was inferred; Pop=1 indicates that a foundress population of 1 foundress per colony was inferred.

